# 3-Mercaptopyruvate Sulfurtransferase (MPST) Regulates Mitochondrial Metabolism and Epithelial Differentiation in Neonatal Patient-derived Airway Cells

**DOI:** 10.64898/2025.12.18.695204

**Authors:** Abhrajit Ganguly, Cynthia M. Carter, Aristides Rivera Negron, Paul T. Pierce, Lynette K. Rogers, Matthew S. Walters, Y.S. Prakash, Trent E. Tipple, Arlan Richardson

**Author notes:** CORRESPONDENCE TO: Abhrajit Ganguly, 800 Research Parkway, Office no: 400E, Oklahoma City, Oklahoma, 73104, USA.

## Abstract

Early-life airway epithelial development relies on tightly coordinated mitochondrial metabolic programs, yet the pathways that govern normal epithelial maturation during this vulnerable developmental window remain poorly defined. Hyperoxia disrupts airway epithelial maturation, contributing to lung injury and airway remodeling in infants with bronchopulmonary dysplasia (BPD), underscoring the need to identify mitochondrial pathways that regulate early epithelial differentiation. 3-Mercaptopyruvate sulfurtransferase (MPST), a mitochondrial sulfur metabolism enzyme, supports mitochondrial metabolic and bioenergetic function, but its role in human airway epithelial development is unknown. In this study, we used neonatal patient-derived tracheal airway epithelial cells (nTAECs) in a three-dimensional air-liquid interface (ALI) model to show that hyperoxia reduces MPST protein abundance. To determine how MPST loss alters early epithelial differentiation and metabolic homeostasis we used RNAi to knock down MPST during ALI differentiation. MPST loss in nTAECs induced early (ALI day 3) transcriptomic shifts involving mitochondrial metabolic pathways, epithelial differentiation programs, and stress-response signature corresponding with decreased ciliated cell numbers during mid-differentiation phase (ALI day 7). Metabolic flux analysis revealed significantly reduced mitochondrial respiration without compensatory increase in glycolysis, indicative of disrupted metabolic flexibility. Together, these data show that MPST is essential for maintaining mitochondrial metabolic integrity necessary for normal airway epithelial development. Loss of MPST creates a developmental vulnerability that may contribute to hyperoxia-induced airway injury in neonates. Targeting MPST-dependent pathways could represent a new strategy to preserve airway health in infants at risk for BPD airway remodeling.

**NEW & NOTEWORTHY:** This study identifies MPST as a previously unrecognized regulator of neonatal airway epithelial development. In our neonatal patient-derived three-dimensional organotypic model, hyperoxia reduces MPST, and MPST loss alters mitochondrial metabolism and epithelial differentiation programs. These findings indicate that MPST deficiency contributes to mitochondrial dysfunction under hyperoxic conditions and highlight MPST-linked pathways as potential therapeutic targets to mitigate early-life airway injury and remodeling relevant to infants with bronchopulmonary dysplasia.

## INTRODUCTION

Airway epithelial developmental programming in early life is highly vulnerable to oxidative stress (1–3). Dysregulated epithelial maturation directly contributes to the abnormal airway remodeling seen in infants with bronchopulmonary dysplasia (BPD) (4–8), a chronic lung disease seen in premature babies. Recent work has highlighted that supraphysiological therapeutic oxygen (hyperoxia) commonly used in Neonatal Intensive Care Units (NICU), provokes broad metabolic reprogramming in neonatal tissues, including the lung, which is driven largely by mitochondrial dysfunction (9). These metabolic shifts are increasingly recognized as early determinants of hyperoxia-induced injury, preceding structural and inflammatory changes. Proper mucociliary development requires timely metabolic shifts regulated by well-functioning mitochondrial bioenergetics (10–12). Recent work in human neonatal airway epithelium shows that even moderately preterm infants (28-29 weeks’ gestation) exhibit diminished mitochondrial function accompanied by defective ciliary development (13), features also reproduced in hyperoxia-based human and mouse models (14, 15). The mitochondrial metabolic programs that govern early airway epithelial lineage commitment remain poorly defined.

3-Mercaptopyruvate sulfurtransferase (MPST) is a redox-active enzyme localized to the mitochondrial inner membrane (16, 17). MPST serves as a key mitochondrial checkpoint that modulates reactive oxygen species (ROS) flux through the mitochondrial electron transport chain (ETC) and maintains bioenergetic function (17–24). MPST preserves mitochondrial complexes (especially Complex I) from oxidative damage (18, 20) via production of the gasotransmitter hydrogen sulfide (H_2_S), and persulfidation, a redox-sensitive post-translational modification of cysteine residues on proteins (17). Using 3-mercaptopyruvate (3-MP) as substrate, MPST generates pyruvate and H₂S, supporting electron flow and mitochondrial respiration (18, 25). MPST upregulation is well documented in human epithelial hepatic (26), renal (27), oral (28, 29) and lung (30) cancers, and enhances epithelial mitochondrial resilience and tumor progression. In contrast, MPST deficiency in humans has been linked to epithelial and metabolic injury across multiple organ systems, including intestinal epithelium in inflammatory bowel disease (19), adipose tissue in obesity (24), and myocardium in heart failure (20). Notably, the role of MPST in the developing human airway epithelium is unexplored.

To interrogate epithelial oxidative injury in the developing neonatal airway, we have established a three-dimensional (3-D) organotypic patient-derived model of human neonatal tracheal aspirate-derived airway epithelial cells (nTAECs) (31). We recently showed that hyperoxia exposure of nTAECs acutely disrupts mitochondrial bioenergetics, driving a metabolic shift from oxidative phosphorylation (Oxphos) toward glycolysis and impairs mucociliary differentiation during air-liquid interface (ALI) culture (15). Importantly, our unbiased proteomic data demonstrated that hyperoxia decreases MPST protein abundance in differentiating nTAECs, suggesting a potential link between impaired sulfur redox metabolism and hyperoxia-induced epithelial dysfunction. Based on this observation, we hypothesized that MPST loss during early airway epithelial development disrupts mitochondrial bioenergetic and metabolic programs, thereby altering epithelial differentiation trajectory. The goals of this study were to: (a) characterize the neonatal airway epithelial phenotype following MPST loss, (b) define the consequences of MPST loss on mitochondrial bioenergetic function, and (c) identify early transcriptomic changes in the developing airway epithelium following MPST loss.

## MATERIALS AND METHODS

### Isolation and Expansion of Neonatal Tracheal Airway Epithelial Cells (nTAECs)

Tracheal aspirate samples were obtained from infants in the NICU after parental consent, using procedures approved by the Institutional Review Board (IRB) of the University of Oklahoma Health Campus (IRB 14377). Fresh samples were processed to isolate nTAECs, which were subsequently expanded and cryopreserved as previously described (31). For this study, nTAECs from two independent donor cohorts were utilized. The first cohort included residual samples of nTAECs from term infants (≥ 37.0 weeks’ gestation) used in our previous hyperoxia exposure study (15) and utilized for complementary validation of MPST protein abundance by immunoblotting following hyperoxia. The second cohort, comprising of nTAECs from term and late-preterm infants (34.0-36.6 weeks’ gestation), was used for siRNA-mediated MPST knockdown experiments. This study was not powered to detect differences by sex owing to the small sample size. nTAECs were subsequently seeded into a 3-D organotypic ALI model for epithelial differentiation.

### Air-liquid Interface (ALI) model and Hyperoxia exposure

nTAECs were differentiated using a 3-D ALI model with 60% O_2_ exposure as previously described (31). ALI differentiation was continued for 14 days (until ALI day 14). On day 7, ALI inserts assigned in the hyperoxia group were incubated in the TriGas incubator (Thermo Fisher Scientific, MA, USA) under 60% O_2_ exposure and continued for 7 days (hyperoxia group) while control inserts remained at 21% O_2_ (normoxia or room air group). On ALI day 14, nTAECs from control and hyperoxia group were harvested for immunoblotting for MPST.

### siRNA transfection in 3D ALI model

*MPST* knockdown was performed using MPST-targeting siRNA SMARTpool or non-targeting Scr control (Dharmacon Horizon Discovery) and transfection was performed at the time of seeding nTAECs on 3-D ALI culture. The siRNA (final concentration 100 nM/well) and the transfection reagent (DharmaFECT-1, final concentration 0.6 µg/well) were mixed in antibiotic-free BronchiaLife^TM^ Epithelial Airway Medium or BLEAM (LifeLine cell technology, MD, USA), and the mixture was added apically to the cell suspension (final apical volume 100 µL). After 24 hrs, both chambers were refreshed with BLEAM (containing antibiotics), and cells were maintained submerged for an additional 24 hrs after which media from the apical chamber was removed, and the basolateral chamber was switched to differentiation media (LifeLine cell technology, MD, USA) to establish ALI (ALI day 0). Because of the transient nature of siRNA-mediated knockdown, 3-D cultures were maintained only through ALI day 7, with media exchanged every 48 hrs. Gene knockdown was confirmed with unbiased transcriptomic analysis utilizing bulk RNA sequencing performed on ALI day 3.

### Immunoblotting

For immunoblotting, nTAECs were harvested from control and hyperoxia groups on ALI day 14 and from siRNA and Scr control groups on ALI day 7. Total protein was extracted directly from ALI wells using RIPA buffer supplemented with protease and phosphatase inhibitors, Laemmli sample buffer, and 2.5% 2-mercaptoethanol. Lysates were resolved on 4–20% SDS-PAGE gels and transferred to nitrocellulose membranes. After blocking in 5% milk/TBST for 1 hour, membranes were incubated overnight at 4°C with MPST (NBP1-82617, 1:2000, Novus Biologicals) or NDUFS1 (12444-1-AP, 1:1000, Proteintech). HRP-conjugated secondary antibody (1:2000, Southern Biotech) was applied for 1 hour, and proteins were detected by ECL using a ChemiDoc imaging system. Densitometry was performed with Image Studio software (version 5.2.1, LI-COR, Lincoln, NE) and normalized to α-Tubulin (ab7291, 1:3000, Abcam) as a loading control.

### Immunofluorescent Staining and Quantification of Epithelial Differentiation

Utilizing ALI inserts from both treatment groups (Scr and siRNA), immunofluorescent staining for secretoglobin 1A1 or SCGB1A1 (club cell; 1:200, BioVendor), and acetylated tubulin or ACTUB (ciliated cell; 1:200, Sigma) was performed using previously published protocol (31). Images were acquired on an Olympus BX43 microscope and at least 6 random fields per insert were analyzed in a semi-automated fashion. DAPI-positive nuclei were counted in an automated fashion using stardist plugin with ImageJ (version 2.160/1.54g, NIH). Epithelial marker-positive cells were quantified manually using ImageJ and normalized to total DAPI-positive nuclei.

### Mitochondrial Stress Test with Seahorse

The Seahorse XFp Mito Stress Test (Agilent, Cat# 103010-100) was performed on an Agilent Seahorse XFp Analyzer following the manufacturer’s protocol. Due to technical limitations in performing the assay on ALI inserts, we utilized undifferentiated basal nTAECs in submerged culture for this experiment. nTAECs were maintained under submerged conditions, transfected with either MPST-targeting siRNA SMARTpool or a non-targeting Scr control (Dharmacon, Horizon Discovery), and incubated for 24 hrs before media replacement. For the Seahorse assay, cells from each treatment group were seeded at 30,000 cells per well into XFp cell culture plates. Oxygen consumption rate (OCR) and extracellular acidification rate (ECAR) were measured at baseline and following sequential injections of oligomycin (1.5 µM) to block ATP-synthase (Complex V) and determine ATP-linked respiration; FCCP or Carbonyl cyanide-p-trifluoromethoxyphenylhydrazone (5 µM) to stimulate oxygen consumption to maximal levels, and a combination of rotenone and antimycin A (0.5 µM each) to inhibit electron transport through Complexes I and III allowing measurement of non-mitochondrial respiration.

### RNA Extraction

Total RNA was extracted directly from 3-D cultures on ALI day 3 (Scr and siRNA group) using the PureLink™ RNA Mini Kit (Thermo Scientific, Cat# 12183018A) according to the manufacturer’s instructions. Residual genomic DNA was removed with an on-column DNase treatment using the PureLink™ DNase Set (Thermo Scientific, Cat# 12185-010).

### Bulk RNA sequencing and Downstream Visualization and Analysis

Bulk RNA sequencing was performed on ALI day 3 samples from both groups (Scr vs siRNA) using standard quality control, read trimming, alignment, and differential expression analysis, with full methodological details provided in the Supplementary Methods. Quality control metrics for all RNA-seq samples, including read quality, GC content, and alignment statistics, are summarized in Supplementary Table S1.

Normalized gene expression data was used for downstream visualization and analysis. A volcano plot was generated using the EnhancedVolcano package to identify genes with both statistically significant and biologically relevant (p < 0.05, |LogFC| ≥ 1) differential expression. For heatmap visualization, the top 20 upregulated and top 20 downregulated genes were selected based on LogFC and plotted using the pheatmap (32) package. Expression values were z-score normalized across samples, and samples were grouped by treatment condition (Scr vs siRNA). For gene set enrichment analysis (GSEA), genes were ranked by LogFC (Scr vs siRNA), and the ranked list was tested against the MSigDB (32) Hallmark gene sets for Homo sapiens obtained via the msigdbr R package. Enrichment was performed with the fgsea algorithm, yielding normalized enrichment scores (NES), nominal p values, and Benjamini-Hochberg-adjusted FDR q values (padj). Hallmark pathways with padj or FDR < 0.05 were considered significantly enriched.

### Statistical methods

Statistical analyses were performed using GraphPad Prism 10.5.0 (GraphPad Software, CA, USA). Fold changes were calculated relative to Scr controls. Because of the small sample size, non-parametric tests were used as noted in the figure legends. A p-value ≤ 0.05 was considered statistically significant. Analysis for bulk RNA-seq dataset is detailed above in the corresponding section. All key genes mentioned in the study including their full names, and associated pathways are summarized in Supplementary Table S2.

## RESULTS

### Hyperoxia exposure of nTAECs acutely downregulates MPST expression in 3D ALI model

We previously demonstrated that hyperoxia exposure during mid-phase of differentiation (ALI day 7 to 14) **[Fig 1A]** disrupts mucociliary differentiation (15) and identified reduced MPST abundance following hyperoxia at ALI day 14 by unbiased proteomics. In the current study, we re-analyzed residual ALI day 14 samples from control and hyperoxia groups by immunoblotting, which confirmed the proteomic finding and demonstrated a significant reduction in MPST (∼30% reduction, p = 0.028) protein across nTAEC donors in response to hyperoxia **[Fig 1B]**.

**Figure 1:**
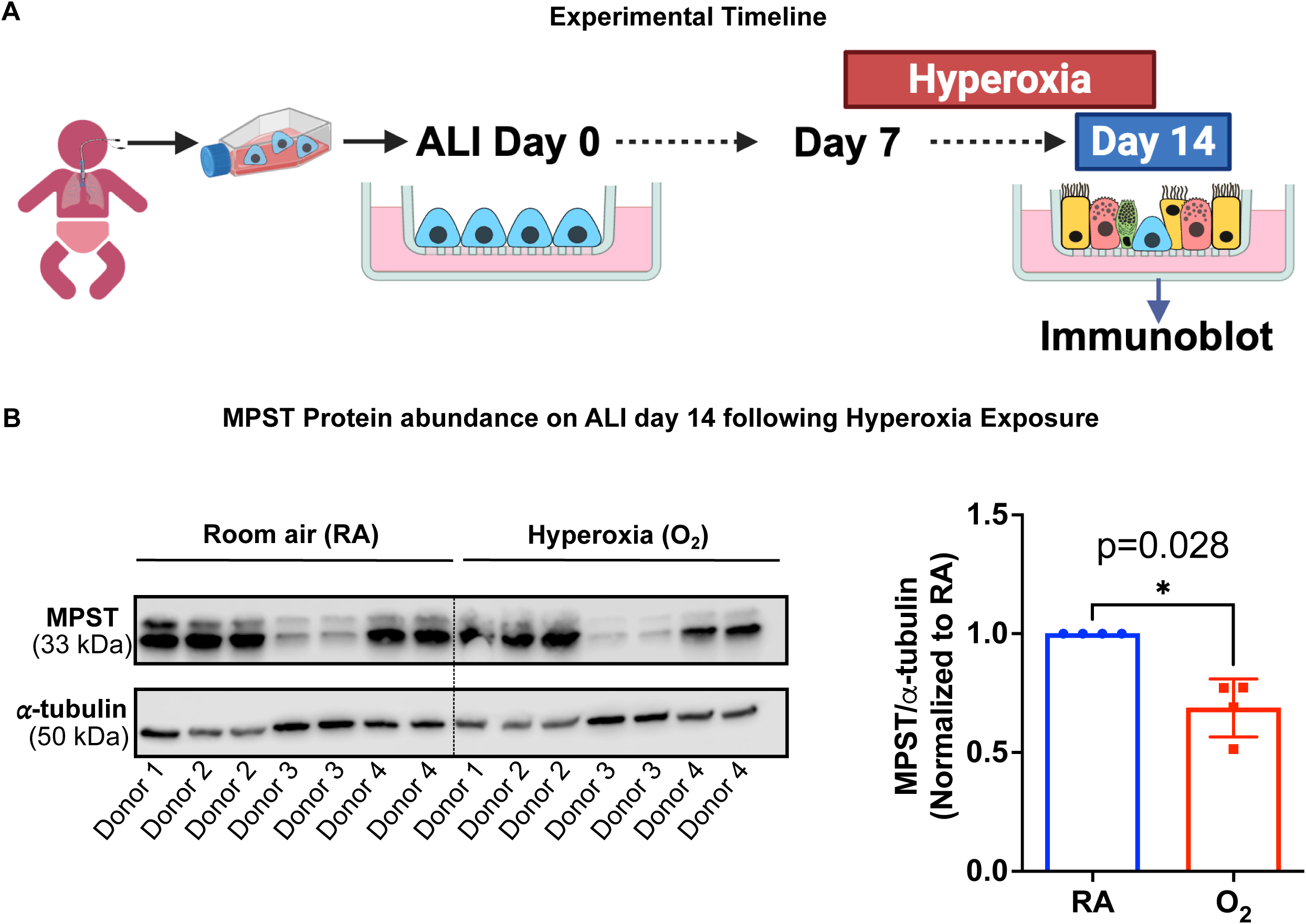
Hyperoxia decreases MPST protein abundance in nTAECs. (A) Experimental timeline for ALI differentiation and hyperoxia exposure (60% O_2_ for 7 days from ALI day 7 to 14) in nTAECs. (B) Immunoblot for MPST on ALI day 14 following hyperoxia exposure of nTAECs. Data points represent protein expression normalized to room air control for each donor. α-Tubulin was used as endogenous control. Statistical analysis was performed using Mann-Whitney U test. Donor numbers (1 to 4) represent corresponding donors in room air and hyperoxia group, n = 4 donor cells, data presented as mean ± sem.

### MPST knockdown of nTAECs alters epithelial differentiation trajectory

To test the hypothesis that MPST loss in nTAECs impairs epithelial differentiation, we induced siRNA-based *MPST* knockdown on nTAECs during early ALI differentiation **[Fig 2A]**. Given the transient nature of siRNA-mediated knockdown, ALI differentiation was continued only through day 7, and ALI day 3 was selected to capture early transcriptomic changes following MPST loss using bulk RNA sequencing. Transcriptomic analysis on ALI day 3 showed effective *MPST* knockdown (∼77% reduction in gene expression, p = 0.007) across donors following siRNA transfection **[Fig 2B]**. We assessed epithelial differentiation on ALI day 7 using immunofluorescent staining of nTAECs with lineage-specific epithelial markers for club (SCGB1A1+) and ciliated (ACTUB+) cells and compared *MPST* siRNA and Scr control groups. On ALI day 7, the siRNA group had significantly decreased ACTUB+ ciliated cells (∼60% reduction, p = 0.007) **[Fig 2C]**. We observed a slight increase (∼15%) in SCGB1A1+ club cells in the siRNA group however, it did not reach statistical significance (p = 0.127).

**Figure 2:**
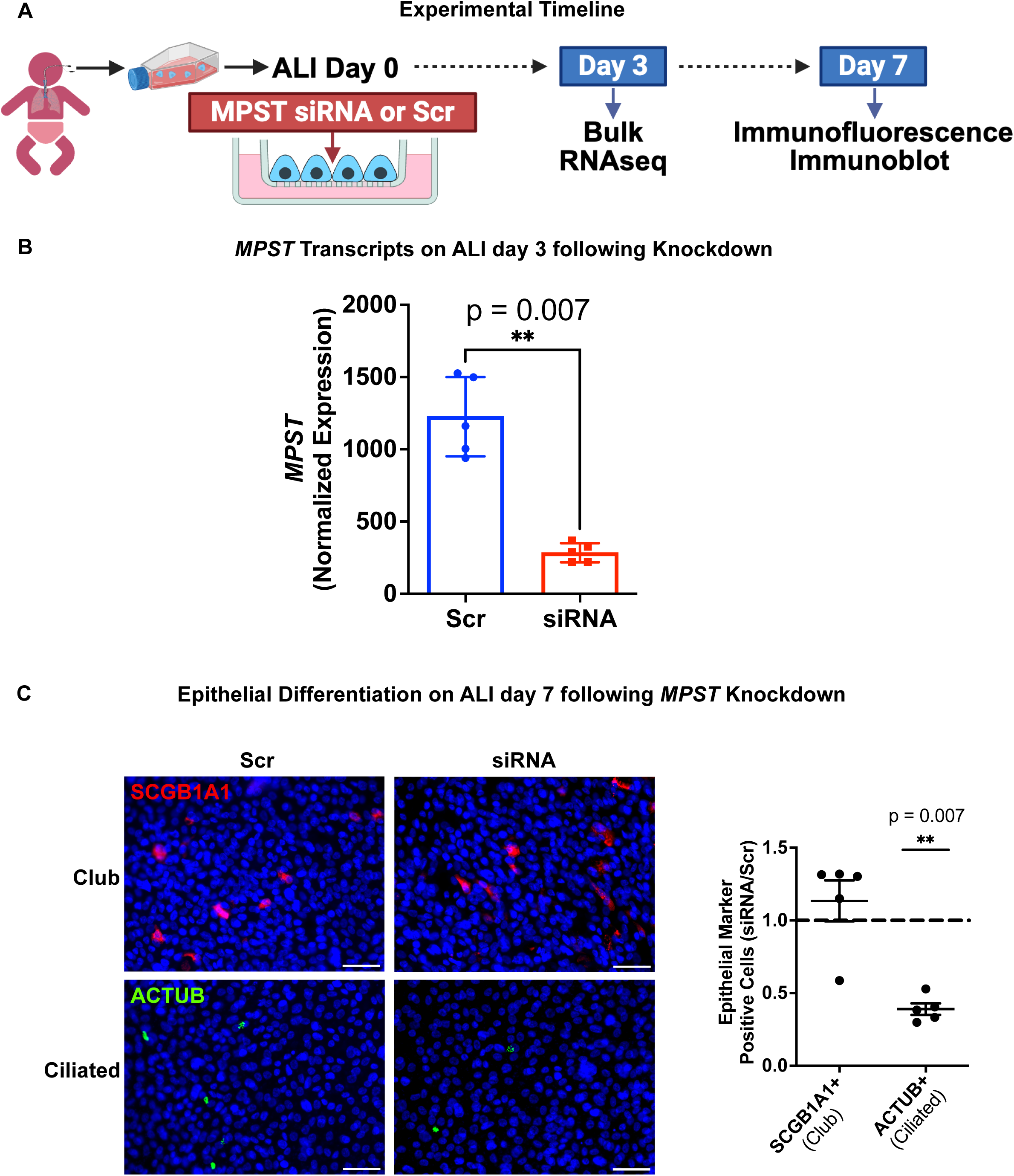
MPST loss alters epithelial differentiation of nTAECs in 3D ALI model. (A) Experimental timeline for *MPST* knockdown during ALI differentiation of nTAECs. (B) Transcriptomic analysis on ALI day 3 showing *MPST* gene expression data following siRNA-based knockdown. Statistical analysis was performed using Mann-Whitney U test, n = 5 donor cells, data presented as mean ± sem. (C) Epithelial differentiation on ALI day 7 using immunofluorescent staining of nTAECs with lineage-specific epithelial markers for club (SCGB1A1+) and ciliated (ACTUB+) cells with the nucleus counterstained by DAPI. The graph showing the positive cells shown for the siRNA treated cells normalized to the Scr control cells for each donor. Statistical analysis was performed using Mann-Whitney U test, n = 5 donor cells, data presented as mean ± sem.

### MPST knockdown reduces mitochondrial respiratory capacity without inducing compensatory increase in glycolysis

Cellular differentiation especially energy intensive processes such as ciliogenesis require intact mitochondrial bioenergetic function (10, 12, 33). Given MPST’s role in cellular energetics, we tested the hypothesis that MPST loss in nTAECs impairs mitochondrial bioenergetic function. We first performed immunoblotting on ALI day 7 and found that *MPST* knockdown reduces NDUFS1 protein abundance (∼35% reduction, p = 0.038), a critical subunit of mitochondrial Complex I and essential for normal bioenergetic function **[Fig 3A]**. To directly assess mitochondrial functional consequence of *MPST* knockdown in nTAECs, we employed Seahorse metabolic flux analysis with Mito Stress Test. MPST knockdown markedly impaired mitochondrial respiration. Basal respiration was significantly reduced in siRNA group (∼27% reduction, p = 0.003), and maximal respiration following FCCP was also significantly diminished (∼43% reduction, p = 0.003) **[Fig. 3B, D]**. Oligomycin-sensitive (∼32% reduction, p = 0.003) and non-mitochondrial respiration (∼33% reduction, p = 0.008) similarly showed significant decreases. ECAR measurements showed downward trends across basal glycolysis (∼14% reduction), oligomycin-stimulated glycolysis (∼19% reduction), FCCP-stimulated glycolysis (∼20% reduction), and non-glycolytic acidification (∼22% reduction), none of which reached statistical significance (all p > 0.24) **[Fig. 3C, E]**.

**Figure 3:**
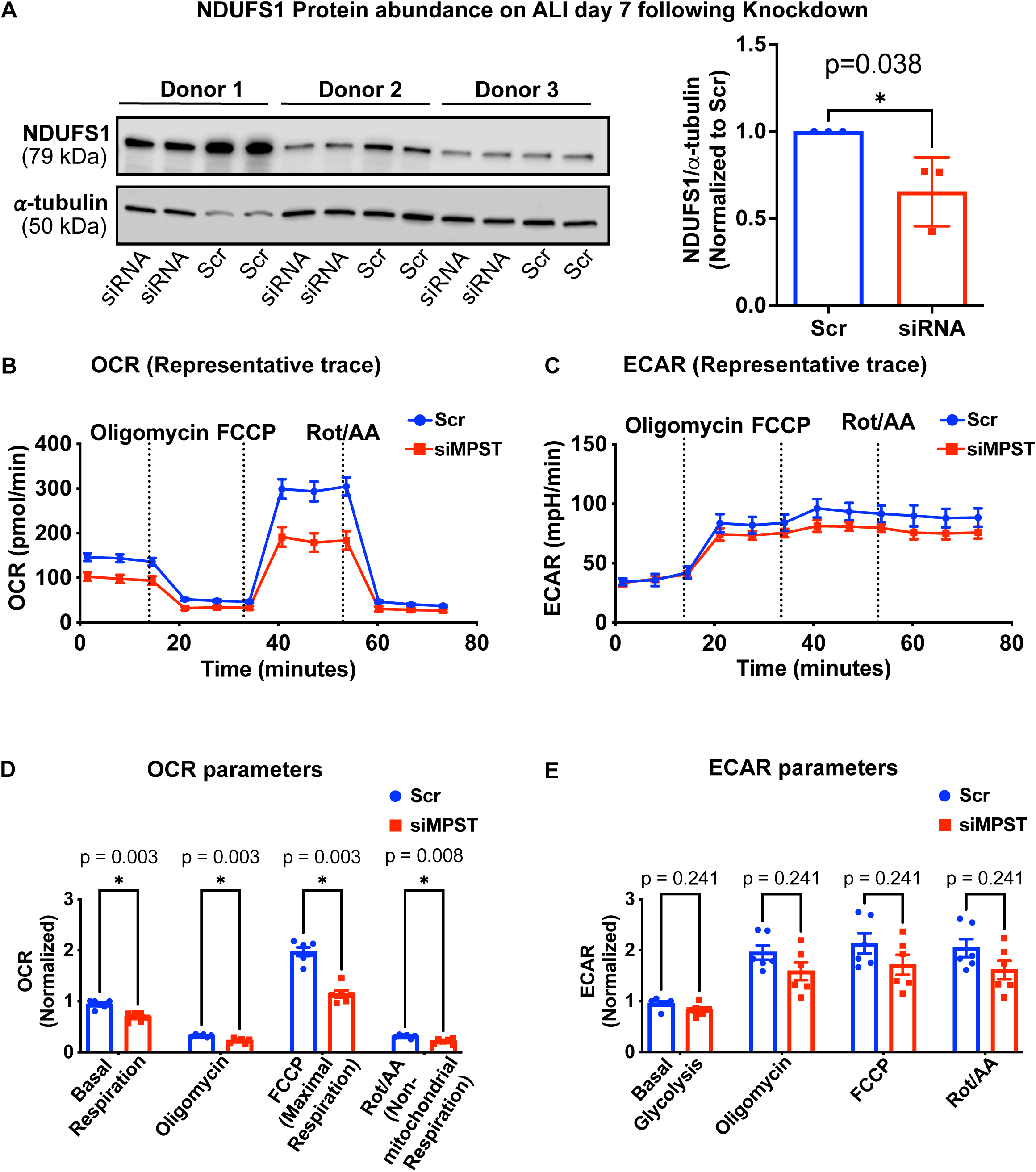
*MPST* knockdown impairs mitochondrial metabolism in nTAECs. (A) Immunoblot for NDUFS1 (critical subunit for Complex I and bioenergetic function) on ALI day 7 following *MPST* knockdown of nTAECs. Data points represent protein expression normalized to Scr control for each donor. α-Tubulin was used as endogenous control. Statistical analysis was performed utilizing Mann-Whitney U test. Donor numbers (1 to 3) represent corresponding donors in siRNA and Scr group, n = 3 donor cells, data presented as mean ± sem. (B - E) Representative traces of oxygen consumption rate (OCR) and extracellular acidification rate (ECAR) from Seahorse Mito Stress Test comparing Scr control and siRNA treated nTAECs across sequential injections of oligomycin (ATP-synthase inhibitor), FCCP (uncoupler), and rotenone/antimycin A (complex I/III inhibitors). Quantified parameters included basal, ATP-linked, maximal, and non-mitochondrial respiration for OCR, and basal and inhibitor-associated glycolytic responses for ECAR. Statistical analysis was performed utilizing Mann-Whitney U test, n = 3 donor cells in duplicates, data presented as mean ± sem.

### Transcriptomic profiling of nTAECs after MPST loss reveals impairment in early epithelial differentiation alongside shifts in metabolic pathways

To determine how *MPST* loss alters pathways linking mitochondrial metabolic pathways and epithelial differentiation, we performed bulk RNA sequencing on ALI day 3. Downstream analysis was performed with assessment of differentially expressed genes (DEGs) following *MPST* loss. We identified 14,974 unique genes and *MPST* knockdown resulted in significant downregulation of 57 and upregulation of 68 DEGs, respectively **[Fig 4A]**. We also plotted the top up and downregulated genes based on changes in log2 fold change. *SCGB1A1* (canonical club cell marker) was one of the top downregulated genes in the siRNA group **[Fig 4B]**, suggesting early disruption in the epithelial differentiation trajectory following *MPST* loss. Additional alterations indicated dampened cytokine signaling (*CXCL5, CSF3*), impairment in Lipid metabolism (*FADS2, PCSK9, ATP10A*) alongside diminished proteostasis regulation (*UCHL1*). Conversely, upregulated genes following MPST knockdown **[Fig 4C]** suggested activation of stress-responsive programs, with induction of interferon-stimulated genes (*XAF1, IFI44L*), inflammatory lipid pathways (*ALOX5*), and extracellular matrix remodeling (*ECM1*). Upregulation of progenitor-associated transcripts (*SHC2, MEGF10, KIT*) together with multiple long non-coding RNAs reflected a shift toward a heightened stress and injury-responsive epithelial state.

**Figure 4:**
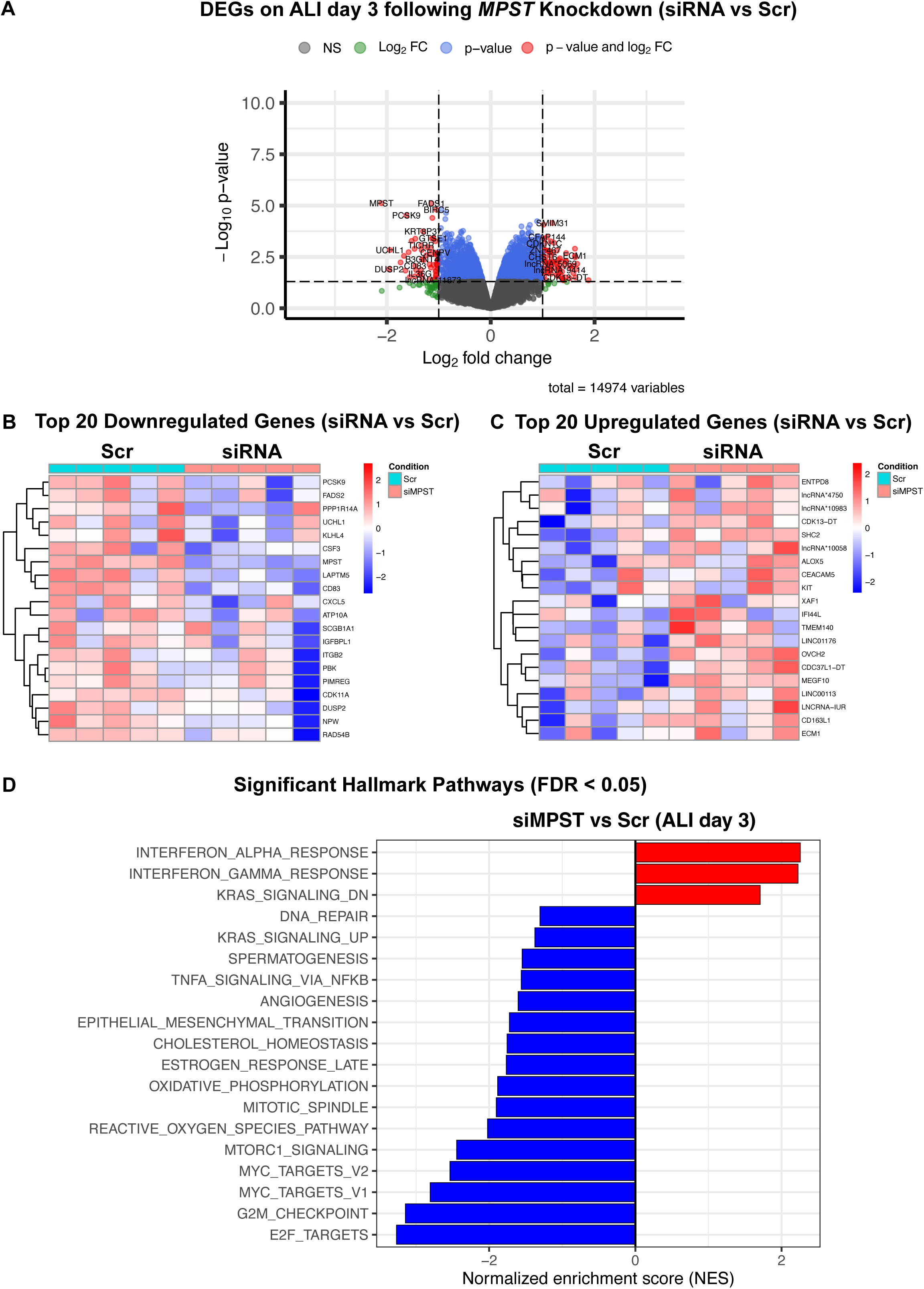
Early transcriptomic changes (ALI Day 3) following *MPST* knockdown in nTAECs. (A) Volcano plot showing differentially expressed genes (DEGs) between siRNA and Scr control nTAECs (n = 5 donor cells). Differential expression was determined using DESeq2 with Wald test statistics and adjusted using the Benjamini-Hochberg method. Red points indicate genes significant by both fold-change and adjusted p-value thresholds; green points denote significant fold-change only; blue points denote significant p-value only. (B-C) Heatmap of the top 20 downregulated and upregulated genes (siRNA vs Scr) following *MPST* knockdown based on log2 fold change. (D) Gene Set Enrichment Analysis (GSEA) using the Hallmark pathway showing significantly (FDR < 0.05) positively and negatively enrichment pathways following MPST loss.

To uncover integrated biological programs that are altered following *MPST* loss in nTAECs, we performed GSEA on the ranked proteome using the Hallmark pathway database **[Fig 4D]**. Negative enrichment was pronounced in Oxphos, mTORC1 signaling, and cholesterol homeostasis, highlighting broad metabolic impairment. The glycolysis pathway also trended toward negative enrichment (NES = −1.29), consistent with our metabolic flux analyses, although this did not reach statistical significance (padj = 0.065). Additional pathways showing significant negative enrichment included cell-cycle and growth programs (E2F targets, G2M checkpoint, MYC targets, DNA repair), epithelial-mesenchymal transition, TNFα-NFκB signaling, and angiogenesis. Positive enrichment was limited to interferon-α and interferon-γ responses and the KRAS signaling (down) set, reflecting targeted activation of interferon-based innate immunity despite global repression of proliferative and metabolic pathways.

## DISCUSSION

The present work demonstrates that MPST is a potential regulator of both epithelial differentiation and metabolic homeostasis in the developing human neonatal airway. Although, MPST is increasingly recognized as a critical mitochondrial transferase that supports redox balance and mitochondrial respiratory chain function (17–24), its role in human neonatal airway epithelial development has been unexplored. By examining early transcriptomic changes following targeted *MPST* knockdown, we show that an early disruption of epithelial differentiation programs (e.g., club and ciliated cell), metabolic pathways and stress-response signatures lead to impaired epithelial maturation trajectory during the mid-differentiation phase. These findings identify MPST as a previously unrecognized node that links mitochondrial metabolism and bioenergetic function with early airway epithelial commitment. The observation that MPST knockdown elicits molecular and bioenergetic signatures overlapping with those induced by hyperoxia suggests that loss of MPST may sensitize the developing airway epithelium to O_2_-mediated injury, thereby contributing to aberrant epithelial differentiation and long-term airway remodeling in infants predisposed to BPD.

This study builds on our prior work (34) showing that moderate hyperoxia exposure (60% O₂, ALI day 7 to 14) drives nTAECs into a persistent progenitor-like state and impairs differentiation into ciliated and goblet cells. We demonstrated that hyperoxia disrupts mitochondrial respiration, downregulates critical Complex I subunits, and induces a metabolic shift with reduced Oxphos and increase in compensatory glycolysis. Our proteomic data also revealed that hyperoxia reduces MPST protein abundance (34). In the current study, we independently confirmed this reduction by re-analyzing residual samples from our earlier work by immunoblotting. To critically test if MPST influences epithelial differentiation we knocked down *MPST* in patient-derived nTAECs during ALI differentiation and evaluated club and ciliated cell differentiation by immunofluorescence at ALI day 7. We observed significant reduction in ACTUB+ ciliated cells following *MPST* knockdown, consistent with an altered epithelial differentiation trajectory. Notably, despite the earlier differentiation timepoint, this phenotype parallels our prior observations at ALI day 14 in hyperoxia-exposed nTAECs, where ciliated cells were similarly diminished (34). Recent studies demonstrate that MPST preserves mitochondrial Complex I integrity and bioenergetic function by limiting oxidative damage (18, 20), and that loss of MPST leads to mitochondrial and metabolic dysfunction across multiple tissues (19, 20, 35). Consistent with this mechanism, immunoblot analysis showed that *MPST* knockdown in nTAECs reduced NDUFS1 protein abundance, a core Complex I subunit which serves as a major entry point for electrons into the ETC and is essential for mitochondrial bioenergetic function (36). This reduction of NDUFS1 mirrors the bioenergetic impairment observed with hyperoxia in nTAECs (34). Collectively, these findings support MPST dysregulation as a mechanistic link between hyperoxia-induced mitochondrial dysfunction and impaired airway epithelial differentiation in the developing lung. Seahorse Mito Stress Test analysis demonstrated that *MPST* knockdown significantly suppressed both OCR and ECAR. In contrast, hyperoxia-exposed nTAECs, despite reduced MPST protein abundance, exhibited a compensatory increase in ECAR, preserving glycolytic reserve (34). Prior studies have shown that hyperoxia induces a shift toward increased glycolytic flux in the lung when mitochondrial respiration is impaired (9, 37, 38). Consistent with this paradigm, hyperoxia-exposed nTAECs retained the capacity for metabolic compensation. In comparison, targeted *MPST* knockdown, which likely resulted in more profound loss of MPST function, abolished this adaptive glycolytic response, indicating a loss of metabolic flexibility. Given the established role of MPST in coordinating mitochondrial electron transport, redox homeostasis, and pyruvate flux (18, 25), significant loss of MPST function likely constrains both mitochondrial respiration and glycolytic capacity. Supporting this concept, in a prior in vivo study (20), MPST deficiency in mouse cardiac tissue resulted in marked mitochondrial abnormalities and metabolic remodeling as revealed by untargeted metabolomic profiling. MPST-deficient hearts exhibited widespread alterations in tricarboxylic acid (TCA) cycle and branched-chain amino acid (BCAA) metabolism, characterized by accumulation of BCAAs and depletion of downstream catabolic intermediates, indicating impaired BCAA catabolism and heightened metabolic inflexibility.

We performed bulk RNA sequencing on ALI day 3 to identify early changes in molecular pathway triggered by MPST loss that precede later epithelial differentiation defects. A key early signature of MPST deficiency was the downregulation of *SCGB1A1*, a canonical club cell marker (39) and an early indicator of mucociliary differentiation. Interestingly, SCGB1A1+ club cells were not significantly reduced at ALI day 7, which may reflect activation of compensatory injury-response programs that club cells typically mount in the airway epithelium (39) or a delayed differentiation trajectory following MPST loss in which early reductions in *SCGB1A1* expression at day 3 partially normalize by day 7. Additional downregulated transcripts reflect broader impairment of epithelial fitness and indicate that *MPST*-deficient nTAECs enter a state marked by diminished capacity for differentiation, lipid remodeling (40), and protein quality control (41, 42) which are essential for lung epithelial maturation. The upregulated genes involved in interferon pathway and extracellular matrix signaling are characteristic of epithelial stress responses and are consistent with a heightened injury-vulnerable state. GSEA highlighted a broad suppression of metabolic and proliferative signaling, including reduced activity in cell-cycle and growth pathways. Negative enrichment of Oxphos and other metabolism-related pathways, such as mTOR (43) signaling and cholesterol homeostasis (44) indicate that MPST loss disrupts cellular metabolic integrity. In a prior study (21), partial knockdown of *MPST* in human adipocytes revealed that proteins with reduced persulfidation - a post-translational modification regulated by MPST - were enriched in metabolic pathways involving pyruvate and glucose metabolism, acyl-CoA biosynthesis, TCA cycle, redox regulation, and sulfur metabolism. Notably, these MPST-dependent metabolic pathways converge with the metabolic programs altered in our nTAEC bulk RNA-seq dataset following *MPST* knockdown, supporting a shared role for MPST in maintaining cellular metabolic programs.

This study has several limitations. First, we used transient siRNA-mediated *MPST* knockdown, which captures early transcriptional and metabolic consequences but may not reflect the sustained alterations produced by chronic MPST deficiency. Future work using stable viral knockdown and complementary overexpression approaches will help define long-term effects of MPST modulation on epithelial differentiation and mitochondrial homeostasis. Second, the nTAECs were derived from term and late-preterm infants and may not fully reflect the metabolic vulnerabilities of extremely preterm airway epithelium. Our future studies will include samples from extremely preterm infants, the population at highest risk of developing BPD and long-term airway remodeling. Third, although bulk RNA-seq and Seahorse flux assays provide strong in vitro evidence, in vivo validation in mouse models using airway epithelial-specific *Mpst* knockout are necessary to establish physiological relevance and determine whether *Mpst* modulates airway development and function under postnatal oxidative stress. Finally, although functional assays following *MPST* knockdown show reduced Oxphos without glycolytic compensation, they do not capture other metabolic pathways that may support epithelial energy needs, such as fatty acid oxidation, glutamine metabolism, or other nutrient sources. Additional targeted Seahorse assays and metabolomic profiling will help determine whether these pathways are also affected and provide a more integrated view of the metabolic consequences of MPST loss.

## CONCLUSION

In summary, our findings demonstrate that loss of MPST disrupts early epithelial differentiation programs and mitochondrial metabolic pathways in neonatal patient-derived airway epithelial cells. These early transcriptomic and bioenergetic defects precede defects in the differentiation trajectory, implicating MPST as a critical regulator of airway epithelial development. Modulating MPST-dependent pathways may provide new opportunities to improve airway health in infants vulnerable to BPD airway remodeling.

## SUPPLEMENTAL MATERIALS

Supplementary methods, Table S1 and S2.

**FigShare link**: https://figshare.com/s/b0df606986c6ca71b54d

## ACKNOWLEDGEMENTS

We thank Ms. Erin Bohon and Aprill Shockley, Neonatal Research Nurses at Oklahoma Children’s Hospital, for their support with participant screening, parental consent, and neonatal sample collection and transport. We also thank the OUHC Institutional Research Core Facilities, including the Genomics Division for RNA library preparation and sequencing, and the Bioinformatics Division for bulk RNA-seq data analysis services. Graphical illustrations were created with BioRender. We acknowledge the use of ChatGPT (OpenAI, 2025) for assistance with language refinement. All AI-assisted edits were reviewed and verified by the authors for accuracy.

## FUNDING

This work was supported by the Presbyterian Health Foundation (PHF) and the HEROES-X Pilot Project Program through funding from PHF to A.G. This work is also supported by funding through the NIH (R01HL160570) to Y.S.P and the Department of Veterans Affairs (1IK6BX005238) to A.R.

## CONFLICT OF INTEREST STATEMENT

No conflicts of interest, financial or otherwise, are declared by the authors.

## AUTHOR CONTRIBUTIONS

A.G. and A.R. conceived and designed the experiments; A.G., C.C., A.R.N and P.P. performed experiments; A.G. analyzed data; A.G. and A.R. interpreted results; A.G. prepared figures and drafted the manuscripts; A.G., L.K.R, M.S.W, Y.S.P, T.E.T and A.R. edited and revised manuscript; All the authors read and approved the final version of the manuscript.

## DATA AVAILABILITY

The bulk RNA seq data has been deposited privately in GEO with the data access information provided separately. The corresponding author will provide original data upon a reasonable request.

## Notes

https://figshare.com/s/b0df606986c6ca71b54d

### Competing Interest Statement

The authors have declared no competing interest.

